# Expanding the Repertoire of Associations Between Antibiotic Resistance Genes and Putative Mobile Genetic Element Carriers Using Metagenomic Long Reads

**DOI:** 10.64898/2026.10.27.762393

**Authors:** Daphne Janelyn L. Go, Anish M.S. Shrestha

## Abstract

Antibiotic resistance genes (ARGs) and the mobile genetic elements (MGEs) that mediate their horizontal transfer are central to understanding the mechanisms, evolution, and dissemination of antimicrobial resistance. Although thousands of ARGs have been documented and numerous ARG–MGE associations have been reported, broader questions about the diversity and distribution of these associations remain underexplored. How comprehensive are the current catalogs of ARG–MGE associations? Are certain MGE families preferentially associated with particular ARGs, and vice versa? How are ARG–MGE associations distributed across microbial taxa? Is the environmental abundance of an ARG related to the diversity of its MGE associations? Do ARG–MGE association patterns vary across environments? Here we address these questions by analyzing multiple publicly available long-read metagenomic datasets from various environmental settings, leveraging the ability of long reads to capture ARGs along with their genomic contexts within individual reads. We identified numerous potential ARG–MGE associations not found in existing databases. A few MGE families were putatively associated with nearly all identified ARGs. ARG–MGE associations were unevenly distributed across taxa. ARG abundance was positively associated with both MGE association diversity and the taxonomic spread of those associations. Furthermore, ARG–MGE association patterns varied across environments. Overall, our findings reveal an extensive diversity of ARG–MGE associations that remains largely underrepresented in current knowledge bases.

**Importance:** Antibiotic resistance is a growing global concern, in part because mobile genetic elements facilitate the spread of resistance genes across bacterial populations, taxa, and environments. Characterizing the associations between resistance genes and mobile genetic elements, and assessing their distribution across taxa and environments, can provide insights into the mobilization and dissemination of antibiotic resistance. Although these associations have primarily been investigated using assembled genomes, our findings demonstrate the potential of unassembled long-read metagenomic data to uncover previously uncharacterized gene–element associations. Our results reveal a substantial reservoir of such associations that remains underrepresented in existing knowledge bases.

## 1 Introduction

Antibiotic resistance genes (ARGs) enable bacteria to survive exposure to antibiotics. They work through various mechanisms. For example, *bla*_*NDM-1*_ and *bla*_*KPC*_ confer resistance to carbapenems through antibiotic inactivation[1, 2]; *mcr-1* alters the target site of colistin[3]; and *mecA* encodes an alternative target protein that reduces binding of methicillin [4]. Several databases curate information on ARGs. One prominent resource is Comprehensive Antibiotic Resistance Database (CARD), which currently contains more than 6,400 ARG sequences covering over 30 drug classes and 8 resistance mechanisms [5].

The acquisition and spread of ARGs are often facilitated by mobile genetic elements (MGE) which can move along with their co-localized ARG cargo horizontally between bacterial cells. Intercellular MGEs like plasmids are responsible for moving ARGs between bacterial cells. ARGs are mobilized onto intercellular MGEs via the activity of intracellular MGEs. Among these intracellular MGEs, IS elements are the smallest, typically encoding only a transposase enzyme required for their own movement, and can affect nearby ARG expression [6, 7]. Two IS elements surrounding an ARG can form a composite transposon, enabling ARG mobilization [7]. For example, *bla*_*NDM-1*_ has been reported on ISAba125-flanked Tn125, and *mcr-1* on ISApl1-flanked Tn6330 [8–10]. Unit transposons encode their own transposase enzymes and mobilize ARGs located within their inverted repeats (IRs) [11, 12]. For instance, *bla*_*TEM*_ is frequently associated with Tn1, Tn2, and Tn3 [13]. Integrons capture ARG-containing gene cassettes through an integrase enzyme and, while not independently mobile, can spread when embedded within transposons or plasmids [14, 15]. Clinically relevant integron cassettes include *β*-lactam and aminoglycoside resistance genes [16–19]. A survey of ARGs and their associated MGEs in clinically relevant species is provided in [13].

Studying the taxonomic and environmental distributions of ARG–MGE associations is important for understanding how antimicrobial resistance (AMR) is maintained and disseminated across bacterial populations and environments. Such analyses can identify key resistance reservoirs, high-risk ARG–MGE combinations in clinically relevant bacteria, and ARGs with greater potential for dissemination. Because bacteria and genetic material move among human, animal, and environmental reservoirs, environmental ARG distributions may also provide insight into AMR dissemination patterns. Additionally, ARG–MGE associations have been used to identify novel ARGs based on the hypothesis that ARGs share conserved MGE contexts [20].

Characterizing the distribution of ARG and ARG–MGE associations involves determining which ARGs are present, identifying their bacterial hosts, assessing their associations with MGEs, and examining how these relationships vary across taxa and environments.

Such analyses have mainly been conducted using assembled bacterial whole-genome sequences or short-read metagenomics datasets [21–26]. A recent effort to catalog ARG–MGE associations is ProMGE, which identified approximately 2.4 million MGEs across more than 76,000 bacterial genomes [21]. However, whole-genome sequence data remain limited, as sequencing efforts tend to focus on a relatively small and over-represented set of clinically relevant species. While metagenomic sequencing captures microbial communities across diverse environments, it primarily relies on short reads, limiting its ability to directly resolve ARG–MGE co-localization. As a result, ARG–MGE associations are sometimes inferred from statistically significant correlations between ARG and MGE abundance profiles [24]. Such correlations, however, do not necessarily indicate physical co-localization as ARGs and MGEs are detected and quantified separately [27]. Although metagenomic assembly can reconstruct longer contigs from short reads, accurately recovering the genomic context surrounding ARGs is challenging, and serious limitations in resolving ARG genomic contexts have been reported for metagenomic assemblers [28–30].

Long-read sequencing technologies have become increasingly prevalent in metagenomic studies in recent years [31–39]. Their substantially longer reads preserve genomic context that is often fragmented in short-read data, enabling multiple genetic features to be captured within a single sequencing read. Here we exploit this capability to identify ARGs together with nearby MGEs, allowing their physical association to be assessed directly [33, 40, 41]. This task is helped by the fact that MGE identification does not require recovery of the complete sequence; instead it relies on detecting conserved marker genes that are characteristic of specific MGE types, such as transposases for IS elements, composite transposons, and unit transposons, or integrases for integrons [33, 40, 42]. Reported ARG–MGE distances are often short, with many associations occurring within 10 kb [43, 44]. These distances fall within typical long-read lengths, allowing many ARG–MGE associations to be captured without metagenomic assembly. Furthermore, the greater sequence information provided by long reads can improve taxonomic classification and profiling, supporting the assignment of ARG and ARG–MGE associations to microbial hosts [45]. Together, these advantages provide an opportunity to examine how ARG–MGE associations vary across bacterial taxa and environmental contexts.

Given this advantage of long reads, this study aims to investigate ARG–MGE associations using publicly available long-read metagenomic sequencing data to characterize the landscape of ARG–MGE associations across diverse bacterial hosts and environmental settings. In particular, we consider the following questions:

- Do long-read metagenomic datasets capture more ARG–MGE associations than those reported based on whole-genome sequences?
- Are certain MGE families more frequently associated with ARGs than others, and vice versa?
- How are ARG–MGE associations phylogenetically distributed?
- How does ARG abundance relate to the diversity of MGEs with which it is associated with and the taxonomic breadth of those associations?
- Do ARG–MGE association patterns vary across environments?

We address these questions by (i) developing a computational pipeline to identify co-localized ARG–MGE pairs in metagenomic long reads, determine their taxonomic origins, and assess whether they are plasmid-borne; and (ii) applying it to 40 publicly available Oxford Nanopore datasets spanning eight environmental sources: municipal wastewater (*n* = 6), community sewage (*n* = 5), activated sludge (*n* = 5), wastewater treatment plants (*n* = 5), swine wastewater (*n* = 4), hospital wastewater (*n* = 5), soil (*n* = 5), and river water (*n* = 5) (See Supplementary Material https://doi.org/10.5281/zenodo.23175603). These environmental categories were the most represented in NCBI SRA, as of October 2025, collectively accounting for *>*90% of the available datasets.

## 2 Results and Discussion

### 2.1 We present a computational pipeline to profile ARGs and co-localized MGEs from long-read metagenomics datasets

Fig. 1 illustrates our pipeline, which uses LAST [46] for alignment because it can perform DNA-DNA and DNA-protein alignments, the latter in a frameshift-aware manner accounting for indel errors common in long-read sequencing data.

**Figure 1:**
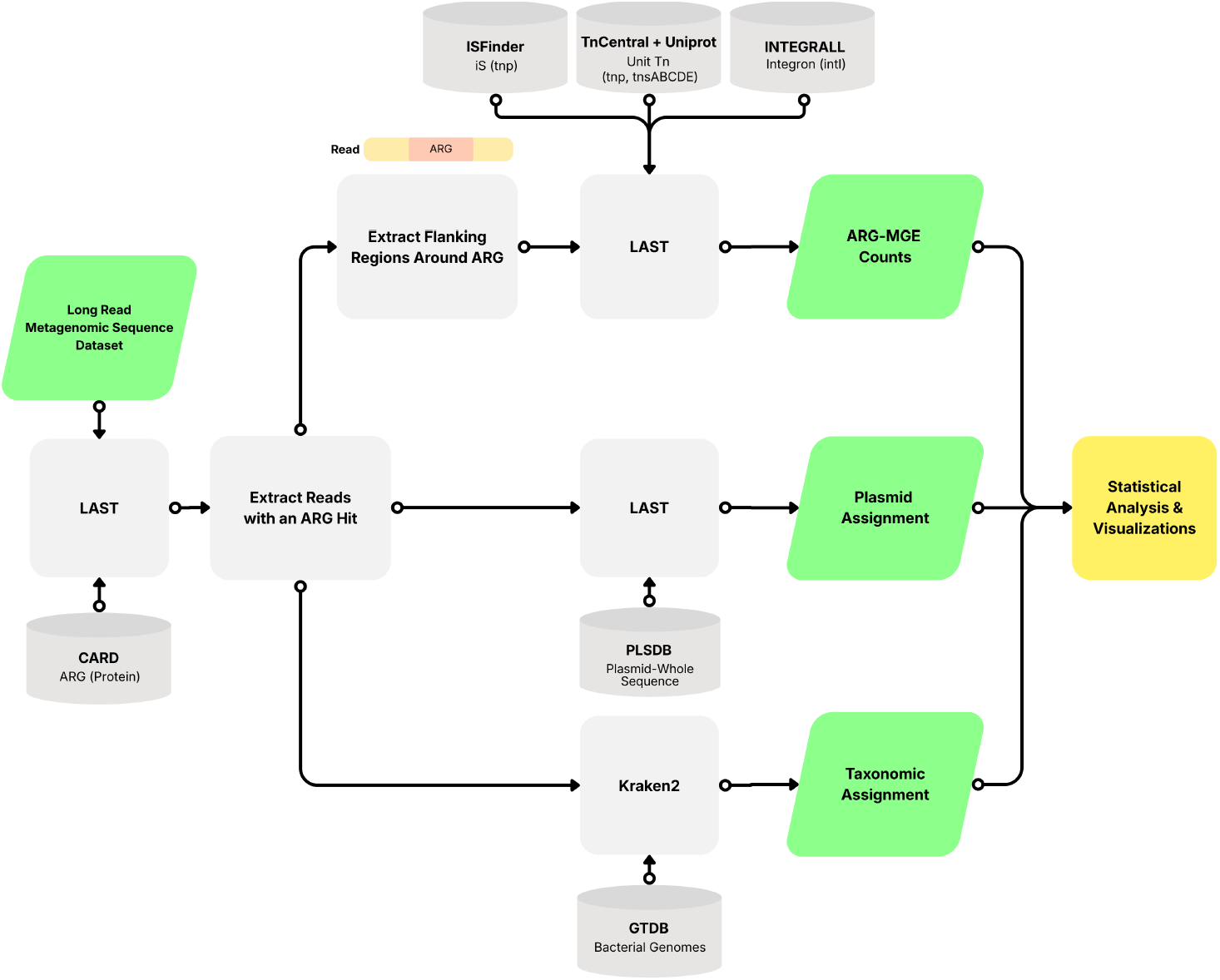
Schematic workflow for ARG and ARG–MGE pair identification, plasmid assignment, and taxonomy classification.

ARGs are identified by aligning reads against the protein homolog model database of the Comprehensive Antibiotic Resistance Database (CARD v.4.0.1 accessed January 2026), which contains more than 6,400 ARG entries, organized into 540 AMR gene families and 46 drug classes using the Antibiotic Resistance Ontology (ARO).

ARG–MGE associations are determined by identifying MGEs co-localized with ARGs on the same read. MGEs are identified by aligning read portions upstream and downstream of ARG hits against specialized MGE databases. IS elements are detected using *>*5,000 transposase sequences across 29 IS families from ISFinder [47](accessed October 2025). Integrons are identified using approximately 1,500 integron-associated *intI* integrase sequences from INTEGRALL [48] (v.1.2, accessed December 2025). Unit transposons from the common Tn3, Tn7, and Tn554 families are identified using transposase and resolvase protein sequences from TnCentral [49] and UniProt [50], with family-specific markers: *tnpA* and *tnpR* for Tn3, *tnsABCDE* for Tn7, and *tnpA, tnpB*, and *tnpC* for Tn554.

Taxonomic classification is performed using Kraken2 [51] against GTDB [52] (release 89.0, accessed February 2026), which contains 145,904 genomes organized into 24,706 species clusters.

Plasmid origin is assessed by aligning reads to PLSDB [53–55] (v.2024 05 31 v2, accessed January 2026), which contains 72,360 plasmid entries.

The pipeline summarizes ARG–MGE co-occurrences, plasmid assignments, and taxonomic profiles for statistical analyses and visualization. Implemented in Snakemake, it requires approximately 35 minutes per Gbp on a machine with 20 CPU cores and 64 GB RAM. The source code is available at: https://github.com/bioinfodlsu/long-read-metagenomics-arg-mge.

### 2.2 We identified many previously uncataloged and unreported ARG–MGE associations

Across approximately 89 million long reads in the dataset, 1% contained an ARG hit. Of these, 4% also contained a co-localized MGE hit. The identified ARG–MGE associations encompassed 1,600 CARD ARG entries spanning 282 ARG families and 60 MGE families, comprising 30 IS families, 26 of which also formed composite transposons, 3 unit transposon families, and integrons. Taxonomic assignment of reads containing ARG–MGE hits spanned 1800 genera. The complete list of identified associations is provided in the Supplementary Material.

We compared the detected ARG–MGE–genus trios with those cataloged in ProMGE [21] (accessed March 2026), a database containing MGEs and their ARG cargo based on a curated subset of assembled prokaryotic genomes from proGenomes2 [56, 57]. To be consistent with proMGE, we grouped the detected IS elements, composite transposons, and unit transposons we detected into a single IS Tn category. ProMGE specifies the ARG cargo of an MGE at the CARD gene family level using the ARO ontology. Accordingly, we aggregated our ARG–MGE–genus trios at the same level to enable comparison.

There is substantial disparity between our set of ARG family–IS Tn–genus trios and those in ProMGE (Fig. 2a). Among the thousands of associations absent from ProMGE, some have been supported by previous reports. Illustrative examples include OXA-48-like *β*-lactamases, *cfr*, nitroimidazole reductase, aminoglycoside bifunctional resistance protein and methicillin-resistant PBP2 whose IS Tn associations are consistent with prior reports [58–63]. Others are potentially novel: FEZ-like, PNGM-family, edeine acetyltransferase, chloramphenicol and rifampin phosphotransferase, and Alm associations with IS Tn elements across multiple genera appear previously unreported. Similarly, Fig. 2b shows a low overlap among our ARG family– integron–genus trios with those in ProMGE. Selected examples of previously reported associations include GIM- and BEL-family *β*-lactamases and aminoglycoside-resistance genes such as *aac(6*^*′*^*)-Ib-cr* and *aadB* [64– 68]. Potentially novel associations include OXA-48-like and OXA-58-like *β*-lactamases with integrons, as well as several glycopeptide-resistance genes, including *vanH, vanX, vanZ, vanU*, and *vanXY* [58, 59, 69–72].

**Figure 2:**
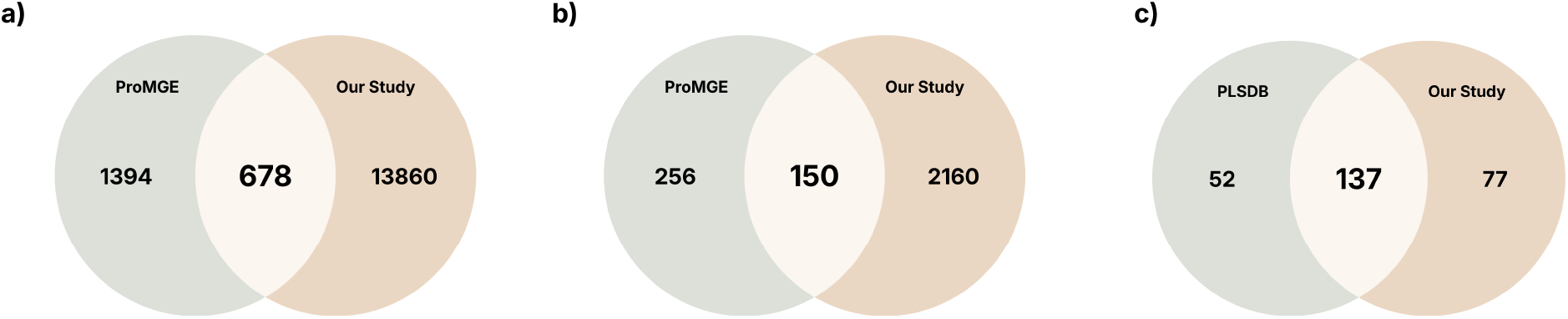
Venn diagrams comparing (a) ARG-family—IS Tn—genus trios we identified with those found in ProMGE, (b) ARG-family—integron—genus trios we identified with those found in ProMGE, and (c) ARG families we classified as plasmid-borne with those annotated in PLSDB.

We also compared the ARGs we classified as plasmid-borne to those annotated in PLSDB [53–55]. Approximately 10% of detected ARG instances were classified as plasmid-borne, lower than previous estimates of 29% in *E. coli* [73] and 31–73% in sewage and activated sludge [33], possibly reflecting our more stringent classification criteria.

Fig. 2c compares our putatively plasmid-borne ARGs, aggregated at the CARD gene family level, with those annotated in PLSDB. We detected 77 ARG families, which were absent from PLSDB. This discrepancy may partly reflect differences in CARD versions used for ARG annotation (v4.0.1 in this study vs. v3.2.9 in PLSDB). To our knowledge, several of these families, including rifampin glycosyltransferase, tunicamycin resistance protein, and VanJ membrane protein families, have not previously been reported on plasmids. Others, including MIR, OXA-427-like *β*-lactamases, and FosC, have been reported on plasmids [74–76], but are absent from PLSDB.

Overall, our findings reveal substantial gaps in current catalogs of ARG–MGE associations, which fail to capture the extensive diversity of ARG–MGE associations present in environmental microbial communities.

### 2.3 A few Tn and IS families putatively carry almost all identified ARGs

We next investigated whether particular ARGs preferentially associate with specific MGEs. For each ARG– MGE pair, we computed the proportion of genera in which the ARG was associated with that MGE, relative to the total number of genera in which the ARG was associated with any MGE type (Eq. 9). The resulting proportions are shown as a heatmap in Fig. 3.

**Figure 3:**
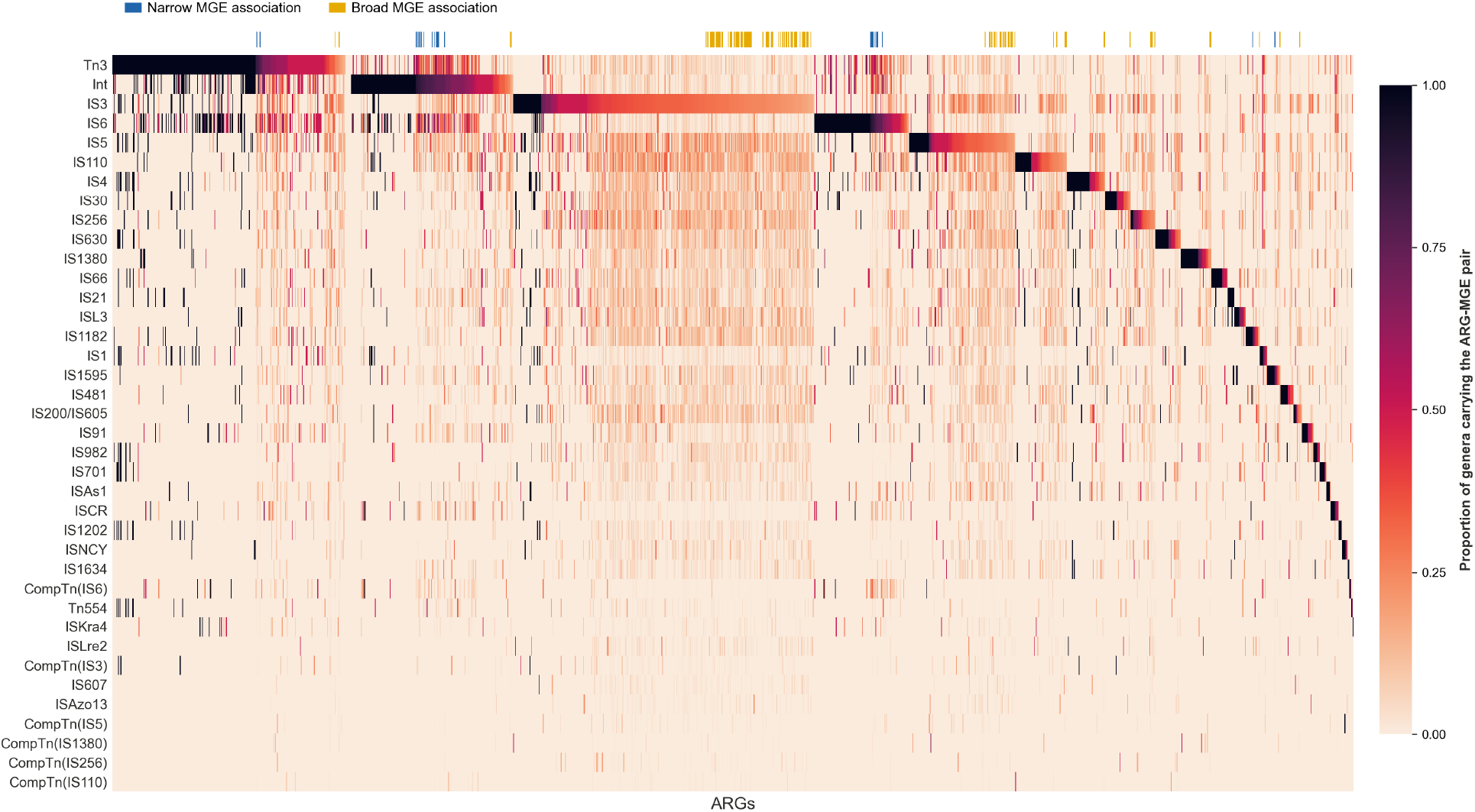
Heatmap showing MGEs (rows) and ARGs (columns), with cell colors indicating the proportion of genera in which the ARG–MGE pair was detected among all genera in which the ARG was associated with any MGE (Eq. 9). ARGs are ordered by their highest-proportion MGE family, and MGE families are ordered by the sum of their proportions across all ARGs. MGEs with proportions below 0.1 for all ARGs and detected in fewer than 2% of genera were excluded from the heatmap. An ARG was classified as having a narrow MGE association if it was detected in ≥ 1% of the genera and one MGE family accounted for a proportion *>*0.7, and a broad association if it was associated with more than 20 MGE families, each accounting for a proportion below 0.3.

Tn3, integrons, IS6, and IS3 were prominent putative carriers of the majority of ARGs, consistent with previous reports linking these MGEs to diverse resistance genes. Tn3-family transposons have been associated with *blaTEM −* 1, *ermAM, strA, strB, tetA(A), van* [77], while integrons commonly capture ARG cassettes such as *aadA, dfrA, cmlA, catB, bla*_*IMP*_, *bla*_*V IM*_, and *bla*_*OXA*_ [78, 79]. Similarly, IS6 and IS3 have been linked to diverse ARGs [80–82], with another study reporting significant associations with ARGs and high abundance in pathogens for IS6, Tn3, IS4, and IS1 [23]. The results could also reflect the fact that if an MGE is common in prokaryotic genomes, it may be more likely to be found near an ARG. This may be particularly relevant for Tn3, and integrons: Tn3-family transposons have been reported across 47 genera and 4 phyla [83], and integrons in approximately 10% of bacterial genomes [84]. Our results suggest that these MGE families are the predominant carriers for a broader repertoire of ARGs than previously documented.

ARGs exhibited varied association patterns. Certain ARGs had a narrow MGE association, predominantly associating with only a few MGE families (columns with blue tick on top row). Examples include *mphA* with IS6, *aadA8* with integrons, and *APH(3”)-Ib* with Tn3. In contrast, some ARGs that associated with a wider range of MGE families (yellow-ticked columns), with no single family predominating: e.g. *RanA, bcrA*, and *tetA(58)*.

### 2.4 ARG abundance is positively associated with MGE diversity and the number of bacterial genera with ARG–MGE associations

Fig. 4 shows the relationship among ARG relative abundance (Eq. 4), the diversity of associated MGEs (Eq. 8), and the taxonomic breadth of the ARG–MGE associations (Eq. 7). The figure suggests a positive relationship between ARG abundance and the number of associated MGE families, with more abundant ARGs generally tending to be associated with more MGE types. ARGs with higher abundance and a greater number of associated MGEs also often occur across more genera, as indicated by larger bubbles.

**Figure 4:**
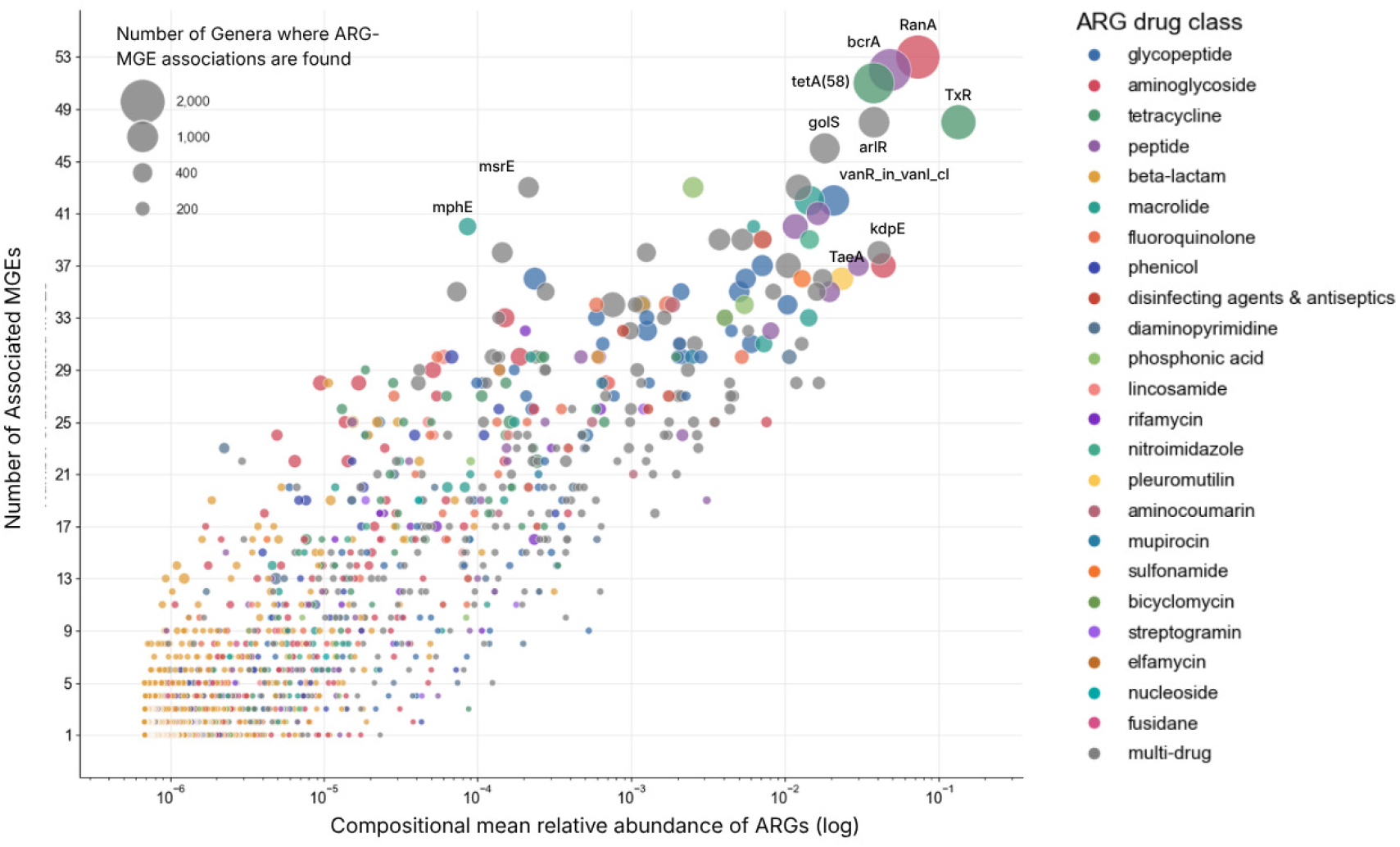
Distribution of ARGs by their mean relative abundance and the number of MGEs they are associated with. The horizontal axis represents the compositional mean relative abundance of an ARG (Eq. 4). The vertical axis represents the number of distinct MGE families associated with an ARG (Eq. 8). Additionally, the bubble size represents the number of genera in which an ARG is found associated with an MGE. The different colors represent the drug classes the ARG is known to confer resistance to.

For several genes in the upper-right of Fig. 4, prior reports provide partial evidence of diverse MGE associations across several genera. For example, *bcrA* has been reported on plasmids and transposon-associated regions in *Enterococcus faecalis* and *Clostridium perfringens* [85–88], while *tetA(58)* has been identified on a mobile plasmid and near MGEs across multiple metagenome-assembled genomes [89, 90]. These reports may represent only a small fraction of a broader, currently underdocumented distribution. Some prominent genes remain poorly characterized with respect to MGE associations. *RanA* and *TxR* have limited evidence of mobility, and although *TxR* has been reported in *Vibrio harveyi* and *Escherichia coli* [91], its MGE associations remain understudied. Likewise, *kdpE* occurs across diverse wastewater and soil metagenomes, but its MGE associations are poorly characterized [92, 93].

Several outliers were also observed. *mphE* and *msrE* had relatively low abundance but were associated with many MGE families. Their association with MGEs is well documented in *Acinetobacter*, including within mobile *pdif* modules and transposon-associated regions [94], but their association with numerous MGE types across diverse genera appears less well characterized.

The bubble colors in Fig. 4 appear to be dispersed across the plot, suggesting that the distribution of ARG–MGE associations is not drug-class specific.

### 2.5 ARG–MGE associations were unevenly distributed across taxa

We next considered the taxonomic distribution of the ARG–MGE associations. To capture association patterns most strongly represented in the dataset, we focused on the 30 predominant ARG–MGE pairs obtained by taking the 15 most abundant ARGs (Eq.4) and for each ARG, the top 2 MGE families they were most frequently associated with (Eq.9). Fig. 5 shows the phylogenetic distribution of the 30 ARG–MGE pairs among the 100 most abundant genera (Eq. 6).

**Figure 5:**
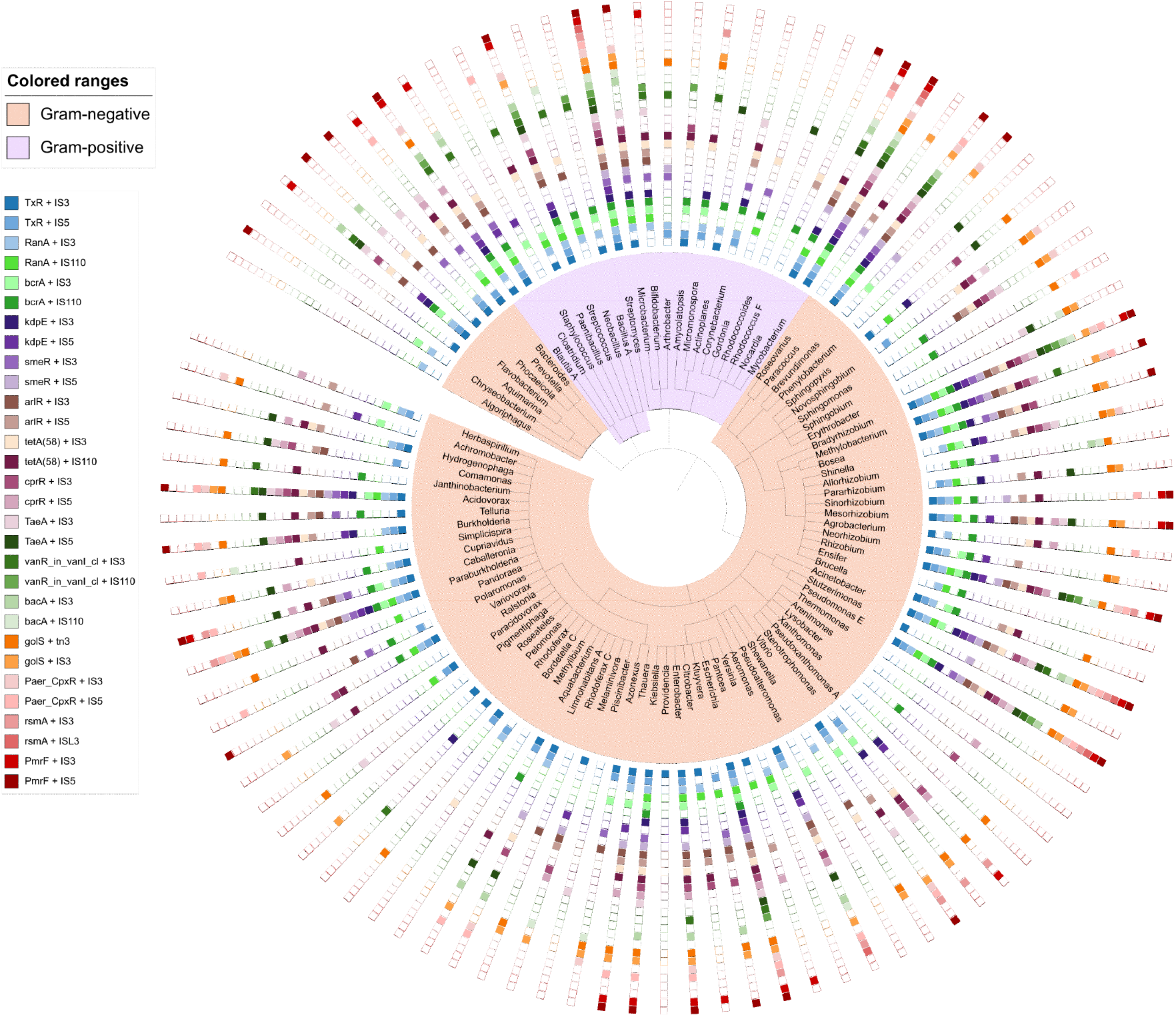
Phylogenetic distribution of the 30 predominant ARG–MGE associations among the 100 most abundant genera (Eq. 6). The 30 pairs were obtained by taking the 15 most abundant ARGs (Eq. 4) and for each ARG, the top 2 MGE families they were most frequently associated with (Eq.9).

We found that ARG–MGE associations were unevenly distributed across taxa, with some genera containing nearly all or all of the 30 dominant ARG–MGE pairs, while others contained few or none. Among the sparsely represented genera were *Algoriphagus, Aquimarina, Providencia, Thermomonas*, and *Staphylococcus*. The limited number of ARG–MGE pairs detected in these genera may suggest that they remain underexplored reservoirs of mobile ARGs. At the other end of the distribution, several genera contained most or all of the selected ARG–MGE pairs, including *Pseudomonas E, Streptomyces*, and *Bradyrhizobium*, potentially reflecting their high species-richness [52]. The presence of potentially mobile resistance determinants across diverse genera highlights the interconnected human, animal, and environmental reservoirs of AMR [95, 96].

Notably, clinically relevant genera such as *Mycobacterium, Escherichia, Acinetobacter*, and *Enterobacter* hosted many of the ARG–MGE pairs. Because our dataset included hospital and urban wastewater, this pattern may partly reflect selective pressures associated with antibiotic exposure [97].

### 2.6 ARG–MGE co-occurrence patterns varied across different environments

Finally, we examined whether ARG–MGE co-occurrence patterns varied across different environments. Fig. 6 shows a PCA based on the relative abundances of ARG–MGE pairs across samples (Eq. 11). Each point represents one sample, while colors indicate the corresponding environment.

**Figure 6:**
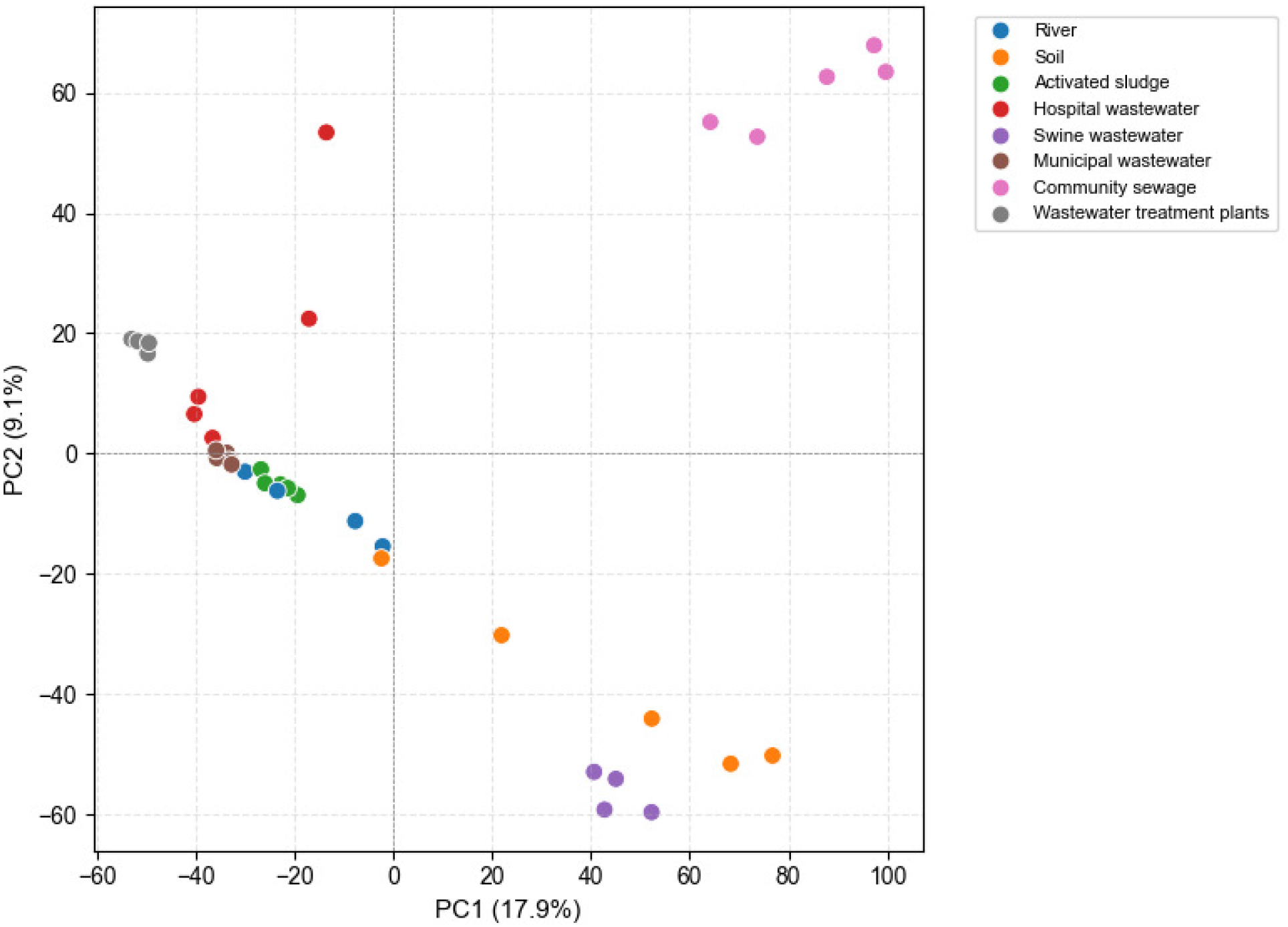
PCA plot of center-log-transformed relative abundance values of ARG–MGE pairs across samples Eq. 11. Each point represents an individual sample and is colored by environmental source.

Clear environment-specific clustering was observed, particularly among community sewage, swine wastewater, and wastewater treatment plant samples.

Previous studies have shown that ARG abundance profiles vary across environments [33, 98]. Differences in ARG abundance and diversity have been reported across environments [98]. Such variation may reflect environmental selective pressures, including antibiotic exposure, while ARG detection in metagenomic data can also be influenced by bacterial diversity, genome size, and gene abundance. The present findings suggest that these factors may also influence ARG–MGE co-occurrence patterns, including their relative abundance and likelihood of detection in metagenomic data.

## 3 Conclusion

We investigated ARG–MGE associations across diverse bacterial taxa and environmental settings using publicly available long-read metagenomic data. By leveraging the genomic context retained within individual reads, we assessed ARG–MGE co-occurrence directly rather than inferring associations from abundance correlations. We identified numerous potentially unreported ARG–MGE associations. Notably, a few transposon and IS families were putative carriers of most detected ARGs. We further found that ARG abundance was positively associated with MGE diversity and the number of genera with ARG–MGE associations. ARG–MGE associations were also unevenly distributed across taxa. Similarly, their co-occurrence patterns varied across environments.

These findings should be interpreted in light of several limitations. First, we defined ARG–MGE associations based on co-occurrence within the same long read, avoiding the need for metagenomic assembly but restricting detection to the length of individual reads. Although this approach is more stringent than proximity-based definitions used in previous studies, which have considered ARGs and MGEs associated when separated by no more than 10 open reading frames [23] or located within 10 kb of each other [99, 100], it may have led us to miss associations spanning longer genomic regions, including nested transposons containing IS elements and integrons. This may also have limited our ability to distinguish individual IS elements from composite transposons. Second, co-localization within a read establishes physical proximity but does not necessarily imply ARG mobility, which would require further functional studies. Moreover, because MGE identification relied on marker proteins rather than complete sequences, some associations may have been overestimated relative to whole-genome-based approaches that consider structural features such as terminal inverted repeats, target site duplications, and recombination sites. Finally, our approach relied heavily on pairwise local alignments, enabling the detection of ARGs and MGE markers even when their full sequences were not captured within a read. However, this may have increased the likelihood of false-positive hits and reduced resolution among closely related gene variants. We therefore grouped genes into CARD families for selected analyses.

Overall, our study demonstrated the use of long-read metagenomics as a complementary approach to surveying ARG–MGE associations across microbial communities, particularly in taxa and environments underrepresented in existing catalogs. The associations identified here provide candidates for further genomic and experimental validation.

## 4 Materials and Methods

### 4.1 ARG Identification and Abundance Quantification

LAST was used to align long-read DNA sequences against CARD using the following commands. The reference database was indexed using `lastdb` with the command `lastdb -q -c CARD-db`. Subsequently, `last-train` was used to estimate alignment parameters, especially the frameshift probabilities, with the command `last-train --codon CARD-db`. Alignments were then performed using the command `lastal -m10 -D1e6 -K1 CARD-db`. Among multiple high-scoring alignments for the same query sequence, we retained only the alignment with the lowest EG value, where EG refers to the expected number of hits between a random query sequence of 1 billion bp and a set of random sequences whose lengths match those in the reference database [46]. We further retained only alignments that covered at least 50% of the reference protein sequence and had an EG value ≤1 *×* 10^*−*5^.

ARG relative abundance was calculated to identify highly abundant ARGs and examine their associations with specific MGE types. For an ARG *A*, its number of copies in a sample [101] was computed using:

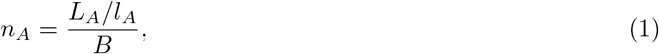

where *L*_*A*_ is the total aligned lengths of reads that hit *A, l*_*A*_ is the nucleotide length of *A*, and *B* is the total number of bases in the sample. The relative abundance of ARG *A* within the sample, denoted by *r*_*A*_, was subsequently calculated as follows:

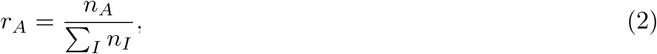

where the summation is across all ARGs *I* detected in the sample.

The mean relative abundance of each ARG across samples was calculated using the compositional mean.

For ARG *A*, let *g*_*A*_ denote the geometric mean relative abundance across samples,

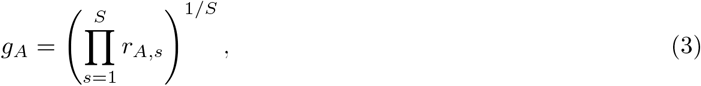

where *r*_*A,s*_ is the relative abundance of ARG *A* in sample *s* and *S* is the total number of samples. The compositional mean *m*_*A*_ was then obtained by closure:

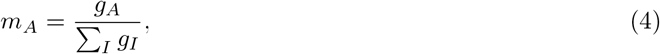

where the sum in the denominator is across all ARGs *I*, so that the resulting values sum to one.

### 4.2 MGE Identification

For each read containing an ARG hit, MGEs co-localized on the same read were identified by aligning the read portions upstream and downstream of the ARG hit to several reference MGE databases. The same LAST alignment procedure used for ARG identification was applied to IS elements, unit transposons, and integrons, differing only in the reference database used. Among multiple high-scoring alignments for the same query sequence, only the alignment with the lowest EG value was retained. MGE hits were required to cover at least 50% of the reference sequence and have an EG value *≤* 1 *×* 10^*−*5^.

IS elements were identified by aligning the extracted regions to transposase sequences from ISFinder [47] (accessed October 2025), which consists of more than 5,000 transposase sequences from 30 IS families. Transposase proteins were used as markers for IS families because they contain conserved active-site motifs required for catalyzing transposition [13, 47, 102].

While individual IS elements do not carry passenger genes, they can mobilize ARGs by forming composite transposons, regions flanked by two IS copies from the same or related family that mobilize as a single unit [13, 49]. Following this, reads containing at least two transposase hits were evaluated against two criteria: (1) the ARG had to be surrounded by transposase hits (i.e., at least one on each side), and (2) the surrounding transposases had to belong to the same IS family. If both conditions were met, the region was classified as a composite transposon.

Integrons were identified by aligning to integron-associated *intI* integrase protein sequences from INTEGRALL [48] (v1.2, accessed December 2025), which contains approximately 1,500 such sequences. The integrase *intI* was used as a marker for integrons because integrons acquire ARGs through gene cassettes that are incorporated through *intI*-catalyzed recombination between the integron’s *attI* site and the cassette’s *attC* site [15, 48].

Unit transposons were identified by aligning to a custom database of protein sequences of transposases and resolvases from the common unit transposon families Tn3, Tn7, and Tn554, obtained from TnCentral [49] and UniProt [50]. Unit transposons were annotated at the family level, with specific transposase and resolvase protein sequences used as markers for each family: *tnpA* and *tnpR* for Tn3, *tnsABCDE* for Tn7, and *tnpA, tnpB*, and *tnpC* for Tn554.

### 4.3 Taxonomy Identification and Quantification

Each read was taxonomically assigned to bacterial taxa using Kraken2 [51, 103] with representative genome sequences from GTDB as the reference database. Representative bacterial genomes with *<*95% completeness or *>*5% contamination were excluded to generate the reference genome dataset. The resulting database was then downsampled using the `--max-db` parameter, with the database size capped at 64 GB. To improve performance for long-read sequence data, both the k-mer and minimizer length parameters in Kraken2 were set to 26 [104]. Relative abundances of microbial species were estimated using Bracken [105]. The Bracken database was built separately for each sample, with the read length parameter set to the average read length of the corresponding sample, using the command `bracken-build -k 26 -l AVG READ LENGTH`.

Genus relative abundances were aggregated across samples using the compositional mean shown below. For genus *G*, let *g*_*G*_ denote the geometric mean relative abundance across samples,

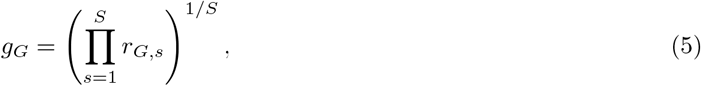

where *r*_*G,s*_ is the relative abundance of genus *G* in sample *s* and *S* is the total number of samples. The compositional mean *m*_*G*_ was then obtained by closure:

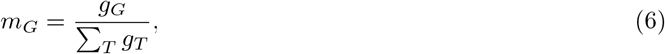

where the sum in the denominator is across all genera *T*, so that the resulting values sum to one.

### 4.4 ARG–MGE Co-occurrence

We say that an ARG *A* is associated with an MGE family *M* in genus *G*, if we find hits for both *A* and *M* on a read that is assigned to genus *G*. We denote the existence of such a read by the indicator variable *I*_*A,M,G*_. Let *S*_*A,M*_ be the set of genera in which the pair *A*-*M* is observed:

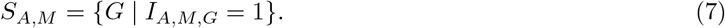

Let *S*_*A*_ be the set of MGE families that *A* is found to be associated with:

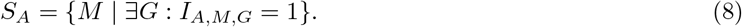

The proportion of *A*’s genus-level association involving *M* is defined as:

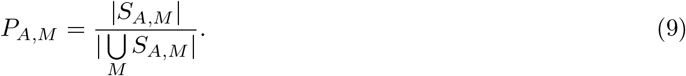

For analysis where we require sample-level abundance of ARG–MGE co-occurrences, we define the following terms. The *A*-*M* co-occurrence count normalized by reference marker lengths and library size is defined as:

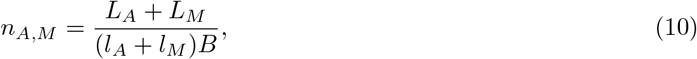

where, from the reads containing hits for both *A* and *M*, *L*_*A*_ is the total aligned lengths of reads that hit *A, L*_*M*_ is the total aligned lengths of reads that hit *M*, *l*_*A*_ is the length of ARG *A, l*_*M*_ is the length of MGE marker *M*, and *B* is the total number of bases in the sample. For MGEs with multiple markers, i.e., Tn3, Tn7 and Tn554, the mean length across all detected gene markers was used. For composite transposons, the length of the bounding IS element is doubled to account for both copies. The relative abundance of pair *A*-*M* within the sample, denoted by *r*_*A,M*_, was calculated as follows:

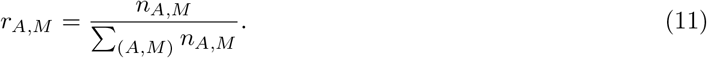

### 4.5 Identifying ARGs on Plasmids

Reads with ARG hits were aligned to whole-plasmid sequences from PLSDB using LAST. The database was indexed with `lastdb plasmid-db`, followed by pairwise alignment with `lastal -m10 -D1e6 -K1 plasmid-db`. For each query, only the alignment with the lowest EG value was retained; alignments with EG *>* 1 *×* 10^*−*5^ or covering *<*90% of the query read were excluded.

## 5 Data Availability

All supplementary data, including the metadata for the publicly available samples, and the complete list of associations identified for each sample are available on Zenodo at https://doi.org/10.5281/zenodo.23492757. The computational pipeline used for data analysis is available at the following URL: https://github.com/bioinfodlsu/long-read-metagenomics-arg-mge.

## 6 Acknowledgments

Daphne Go acknowledges support from the Philippine Department of Science and Technology and the Engineering Research and Development for Technology (ERDT) scholarship program for funding her degree program.

## Notes

https://doi.org/10.5281/zenodo.23492757

### Competing Interest Statement

The authors have declared no competing interest.

